# Spatial Multi-Omics Workflow and Analytical Guidelines for Alzheimer’s Neuropathology

**DOI:** 10.1101/2025.11.16.688710

**Authors:** Xuehan Sun, Hannah R. Hudson, Timothy C. Orr, Srinivas Koutarapu, Alyssa Rosenbloom, Matthew Ingalls, Oliver Braubach, C. Dirk Keene, Joseph M. Beechem, Miranda E. Orr

## Abstract

Spatial biology technologies enable high-dimensional profiling within intact tissues, revealing how molecular and cellular organization drives function and disease. As these platforms gain broader adoption, standardized analytical frameworks are needed to ensure data quality and reproducibility. Here, we present an end-to-end pipeline for the GeoMx Digital Spatial Profiler that simultaneously generates whole-transcriptome and 637-protein measurements from user-defined regions within the same tissue sections. The workflow integrates morphology-guided region selection, quality control, normalization, and multi-modal data interpretation. Applied to formalin-fixed cortical tissues from Alzheimer’s disease, dementia with Lewy bodies, amyotrophic lateral sclerosis, and controls, the framework resolves spatially distinct molecular domains. Transcript and protein signals diverge across amyloid plaque cores and surrounding glial-rich regions, with RNA–protein concordance varying by disease condition, while single-neuron profiling with and without pathogenic tau deposition illustrates protein assay sensitivity. This dataset provides a rigorously validated resource for spatial multi-omic analyses and establishes broadly applicable guidelines for reliable, reproducible profiling of complex tissues.

## Introduction

Spatial organization underlies the function of biological systems, from single cells to complex tissues. Spatial multi-omics technologies now enable high-plex measurement of RNA and protein within intact tissues, linking molecular states to their architectural context. However, standardized analytical frameworks for these datasets remain limited, particularly for integrated transcriptomic-proteomic analyses. Neurodegenerative diseases exemplify the need for such integrative approaches. Alzheimer’s disease (AD)^1^ and related disorders, including dementia with Lewy bodies (DLB)^2^ and amyotrophic lateral sclerosis (ALS)^3^, exhibit spatially heterogeneous pathology that challenges conventional molecular profiling^4–7^. Amyloid plaques are defining, but not exclusive hallmarks of AD: they are also observed in Parkinson’s disease, (DLB)^8,9^, (ALS)^10^, and even in some clinically normal older adults^11^. Although defined as extracellular aggregates of amyloid β (Aβ), a cleavage product of amyloid precursor protein (APP)^12^, plaques contain dozens of associated proteins reflecting complex molecular composition^13,14^. Transcriptional changes in plaque-adjacent tissue are similarly context dependent, varying by disease stage, brain region, and local cell composition^5,15–17^. Because plaque cores are largely acellular and RNA abundance is low, transcriptomic data alone cannot fully resolve their biology. Post-transcriptional regulation and differences in assay sensitivity further decouple RNA and protein signals, underscoring the need for integrated, multiomodal, spatially resolved measurements.

To address these challenges, we applied GeoMx Digital Spatial Profiling (DSP), a high-plex platform that uses oligonucleotide-labeled antibodies and RNA probes to quantify molecular targets within user-defined regions of interest in formalin-fixed, paraffin-embedded human brain tissue^18^. We profiled 637 proteins and the whole transcriptome across amyloid plaque–centered and matched control regions collected from primary motor and visual cortices of non-demented controls (NDC), AD, DLB, and ALS cases. Recognizing that housekeeping genes and proteins vary across disease states and pathologies, we systematically evaluated normalization strategies and analytical approaches to ensure data robustness. Integrated RNA–protein analyses revealed spatially distinct molecular domains and frequent RNA–protein discordance.

We further evaluated the protein assay sensitivity by profiling individual neurons with and without pathogenic tau deposition as neurofibrillary tangles (NFTs), demonstrating single-cell resolution and detection of neuronal heterogeneity. Finally, a subset of GeoMx protein targets was validated using CellScape, a cyclic immunofluorescence platform that directly detects proteins at subcellular resolution. This orthogonal validation reinforces the accuracy and reproducibility of proteomic measurements.

Collectively, these studies establish a biology-informed framework for spatial multi-omics that integrates RNA and protein landscapes with tissue architecture. The resulting dataset serves as a validated resource and practical guide for rigorous, reproducible spatial multi-omic profiling of complex tissues in neurodegeneration and beyond.

## Results

### Study design and conventional baseline Quality Control (QC)

We used four fluorescently labeled morphology markers to guide ROI selection: Syto83 for nuclei, a MAP2 and HuD antibody cocktail for neurons, AT8 (anti-pTauSer^202^/Thr^205^) for neurofibrillary tangles (NFTs), and 6E10 (anti-Aβ N-terminal) for amyloid plaques. These markers were chosen to visualize biologically relevant features without competing with protein panel targets^19^. ROIs were selected from primary motor cortex (PMC) and visual cortex (VC) across four neuropathological diagnoses: NDC, AD, DLB, and ALS (Fig. 1A; Table S1). For plaque-associated ROIs, the Center area of illumination (AOI) encompassed the 6E10-positive regions, Ring1 surrounded the Center, and Ring2 surrounded Ring1 (Figs. 1B–C). Adjacent regions lacking 6E10 immunoreactivity were designated as within-tissue negative controls.

**Figure 1.**
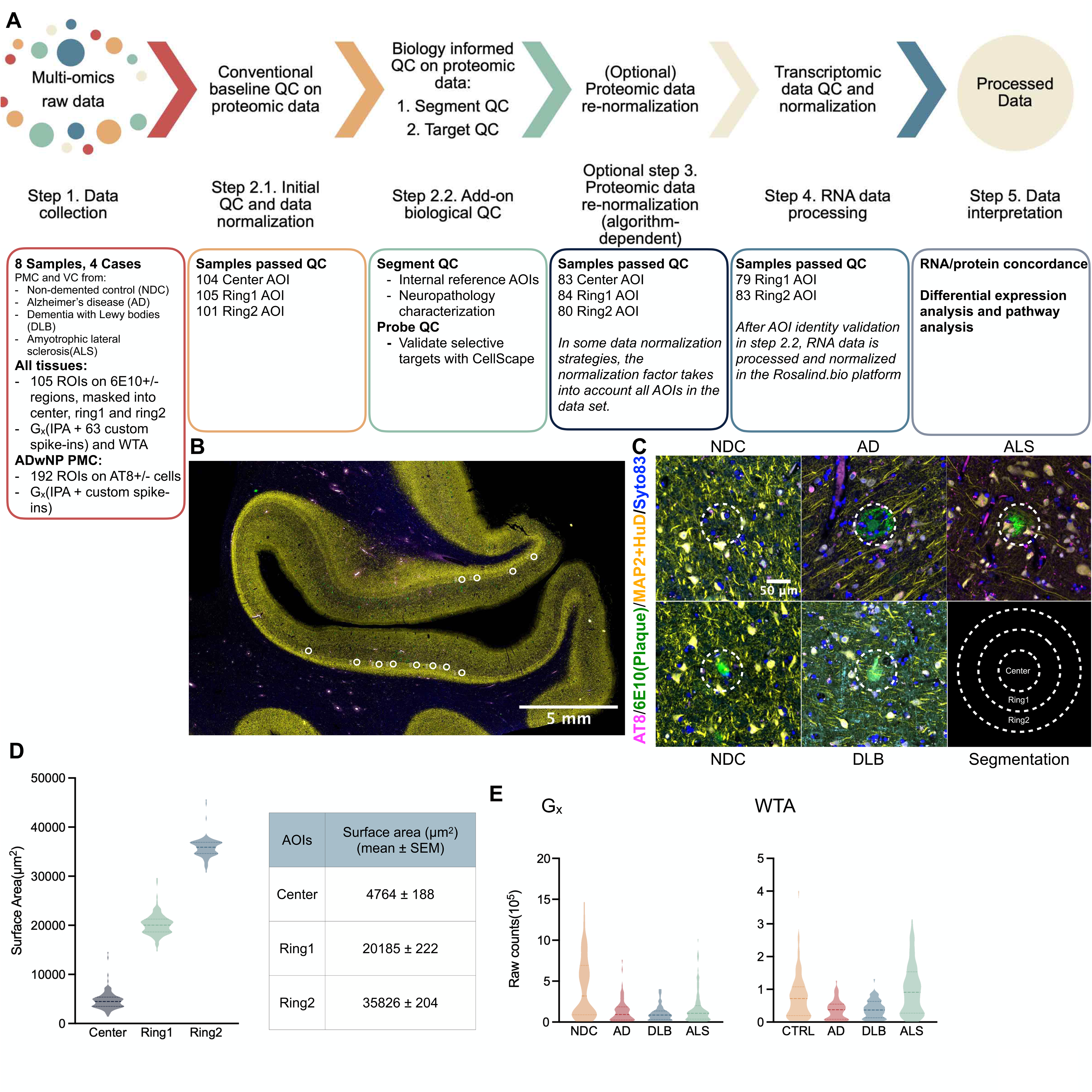
Overview of study design, sample processing, and data analysis. A. Schematic of the study design and data analysis workflow. Primary motor cortices (PMC) and visual cortices (VC) from 4 disease conditions (NDC, AD, DLB, and ALS) were selected for GeoMx DSP multi-omics analysis. ROIs, including plaque and control areas, were selected for both proteomic and transcriptomic profiling (probes used: G_x_, immuno-oncology proteome atlas plus customized 63 spike-in proteins; WTA, whole transcriptome analysis). Each ROI was segmented into 3 AOIs (Center, Ring1, and Ring2) based on 6E10 staining for downstream analysis. B. Representative immunofluorescent staining of tissue sections. ROI selections were highlighted by white circles. C. Representative on-tissue control and 6E10 staining. Plaques were labeled using the 6E10 antibody (green, highlighted by white dashed circles), neurons were stained with MAP2+HuD (yellow), tau phosphorylated at Ser^202^/Thr^205^ was stained with AT8 (magenta), and nuclei were stained with Syto83 (blue). In each condition, various 6E10 staining patterns were detected. The dashed circles in the lower right panel illustrated the segmentation strategy (not to scale). D. Surface area of collected AOIs. E. Raw counts per AOI across disease conditions. Left: G_x_ probe counts; right: WTA probe counts.

6E10-positive stains displayed diverse morphologies and sizes across disease conditions, necessitating flexible AOI dimensions (Fig. 1D). We quantified protein and RNA expression using a curated 637-protein panel (G_x_), which combines the Immuno-oncology Proteome Atlas with 63 spike-in proteins to interrogate neuropathology, immune states, stress responses, and cellular senescence^20^ and Whole Transcriptome Atlas (WTA) consisting of ∼18,000 RNA targets (Fig. 1E).

Proteomic data were processed first, using conventional baseline QC and normalization in the GeoMx DSP Analysis Suite, followed by additional biology-informed QC. This included evaluating amyloid plaque morphology with Cellscape, a cyclic immunofluorescence system with high-dynamic-range microscopy, and confirming 6E10 panel expression; AOIs lacking clear plaque signal were excluded (Fig. 1A).

Across all ROIs, comparable numbers of total cells and neurons were profiled (Figs. S1D–E). Despite similar cell counts, AD and DLB cases exhibited lower protein and RNA detection, including fewer raw reads and detected genes (Figs. 1E, S1A–C). While some group-wise differences were statistically significant, overall variance was small, and cell numbers did not correlate with raw read counts.

### Considerations for proteomic data normalization

Raw protein counts correlated with AOI surface area (e.g., Center < Ring1 < Ring2; Fig. S1A). To enable comparison across AOIs of differing sizes, we applied surface area-based scaling: each AOI’s surface area was divided by the minimum AOI area in the dataset. This adjustment effectively scaled all counts relative to the smallest AOI.

We next evaluated five housekeeping proteins (RPS6, calreticulin, GAPDH, histone H3, TOMM20) and negative control IgGs using the “Evaluate-Normalization-Options” script to assess concordance and variance. Calreticulin, RPS6, and TOMM20 exhibited the highest concordance and lowest variability (Figs. 2A, S2A). All housekeeping proteins displayed disease-dependent variation (Fig. 2B). Since normalization to these proteins could obscure biologically meaningful differences, we did not pursue this strategy in downstream analyses. To assess background non-specific binding in the antibody-based GeoMx panels, we examined negative control IgGs (rabbit, mouse IgG1/2b, hamster, and rat IgG2a; Figs. 2C-E). In GeoMx, IgG controls can be used for normalization by scaling AOIs relative to the dataset-wide geometric mean, or for background correction, which divides each protein’s raw count by the AOI-specific IgG geometric mean. Normalization balances variation across segments, whereas background correction primarily reduces disproportionately high signals.

**Figure 2.**
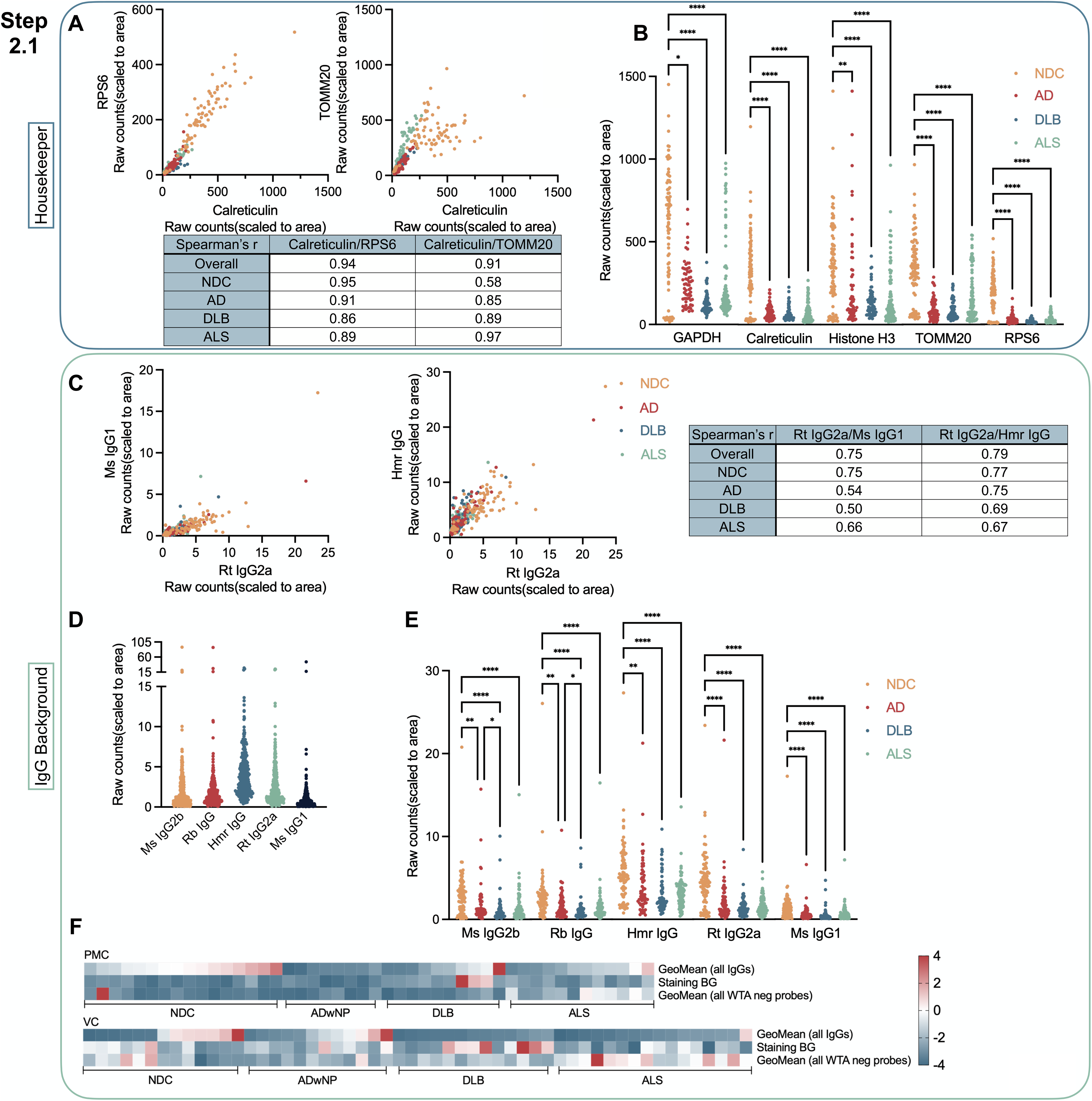
Comparison of normalization strategies for GeoMx protein readout. A. Correlation between calreticulin and two additional housekeeing probes (RPS6 and TOMM20). Each point represents a segmented AOI, with colors indicating disease condition. Probe counts were scaled to the surface area of each AOI to account for size differences. Since the data were not normally distributed, Spearman’s correlation coefficients (r) are reported. B. Expression levels of housekeeping probes across disease conditions. Probe counts were scaled to the surface area of each AOI to account for size differences. Significance was determined using Kruskal-Wallis Anova tests with Dunn’s correction. C. Correlation between Rat IgG2a (used as a negative probe) and two additional negative probes (mouse IgG1 and hamster IgG). Each point represents a segmented AOI, with colors indicating disease condition. Probe counts were scaled to the surface area of each AOI to account for size differences. Since the data were not normally distributed, Spearman’s correlation coefficients (r) are reported. D. Count level of IgGs. Probe counts were scaled to the surface area of each AOI to account for size differences. E. Count level of IgGs across disease conditions. Probe counts were scaled to the surface area of each AOI to account for size differences. Significance was determined using Kruskal-Wallis Anova tests with Dunn’s correction. To better visualize the data, one AOI from the ALS case was excluded from the plot due to its high values (Ms IgG2b = 89.7, Rb IgG = 88.7, Hmr IgG = 24.8, Rt IgG2a = 24.3, Ms IgG1 = 45.8). F. Concordance between IgG counts, staining background, and WTA negative probe counts. For each IgG and WTA negative probe, counts from individual AOIs were summed to obtain values for each ROI. The geometric mean was then calculated across all IgGs or WTA probes. Staining background was calculated by measuring the mean pixel intensity in unstained areas of the 6E10 channel. The three metrics were scaled to a range of -4 to 4 and plotted on the heatmap (see Methods for details).

In our dataset, IgG controls showed high variability and poor concordance, especially at low counts, and elevated IgG signals were limited to a subset of AOIs (Figs. 2C–E, S2B). Comparison with two additional background measures, mean 6E10 fluorescence and negative WTA probe counts, revealed low correlation (Fig. 2F). Notably, elevated IgG counts in the NDC case did not correspond to these measures, indicating that IgGs did not reliably reflect non-specific binding and were unsuitable for normalization.

To assess single-cell proteomics, we extended these methods to single-neuron profiling in AD PMC tissue (25 μm-diameter ROIs around AT8-positive and -negative neurons in cortical layers II and V; Fig. S3E). Housekeeping protein expression varied between AT8-positive and - negative neurons and across cortical layers (Fig. S3A). IgG background signals were inconsistent across ROIs, particularly at low counts, and AT8-negative neurons often displayed more variable IgG signals than AT8-positive neurons (Figs. S3B–F). Because all single-cell ROIs were identical in size, area scaling was inapplicable. However, ROIs with elevated IgG levels also exhibited disproportionately high total target counts, suggesting non-biological signal inflation (Fig. S3F). Therefore, for single neurons, we applied signal-to-background correction, dividing raw counts by the AOI-specific geometric mean of IgG controls.

Finally, we demonstrated that the choice of normalization strategy profoundly affects downstream analyses. In the plaque dataset, proteins identified by differential expression (DE) analysis between 6E10-positive and -negative AOIs varied substantially across three approaches: housekeeping normalization, area scaling, and IgG-based background correction (Fig. 3). Each method identified largely distinct sets of differentially expressed proteins (DEPs), with markedly different distributions in volcano plots (Figs. 3A, 3B). Similar effects were observed in single-neuron ROIs (Figs. 3C, 3D), with housekeeping normalization producing highly biased signals. These results highlight that, in some cases, proceeding with minimally processed or raw data may be the most appropriate approach. Based on these observations, we scaled plaque AOI counts to the surface area of the smallest plaque without applying housekeeping or IgG-based normalization, while choosing signal to background correction for single neurons.

**Figure 3.**
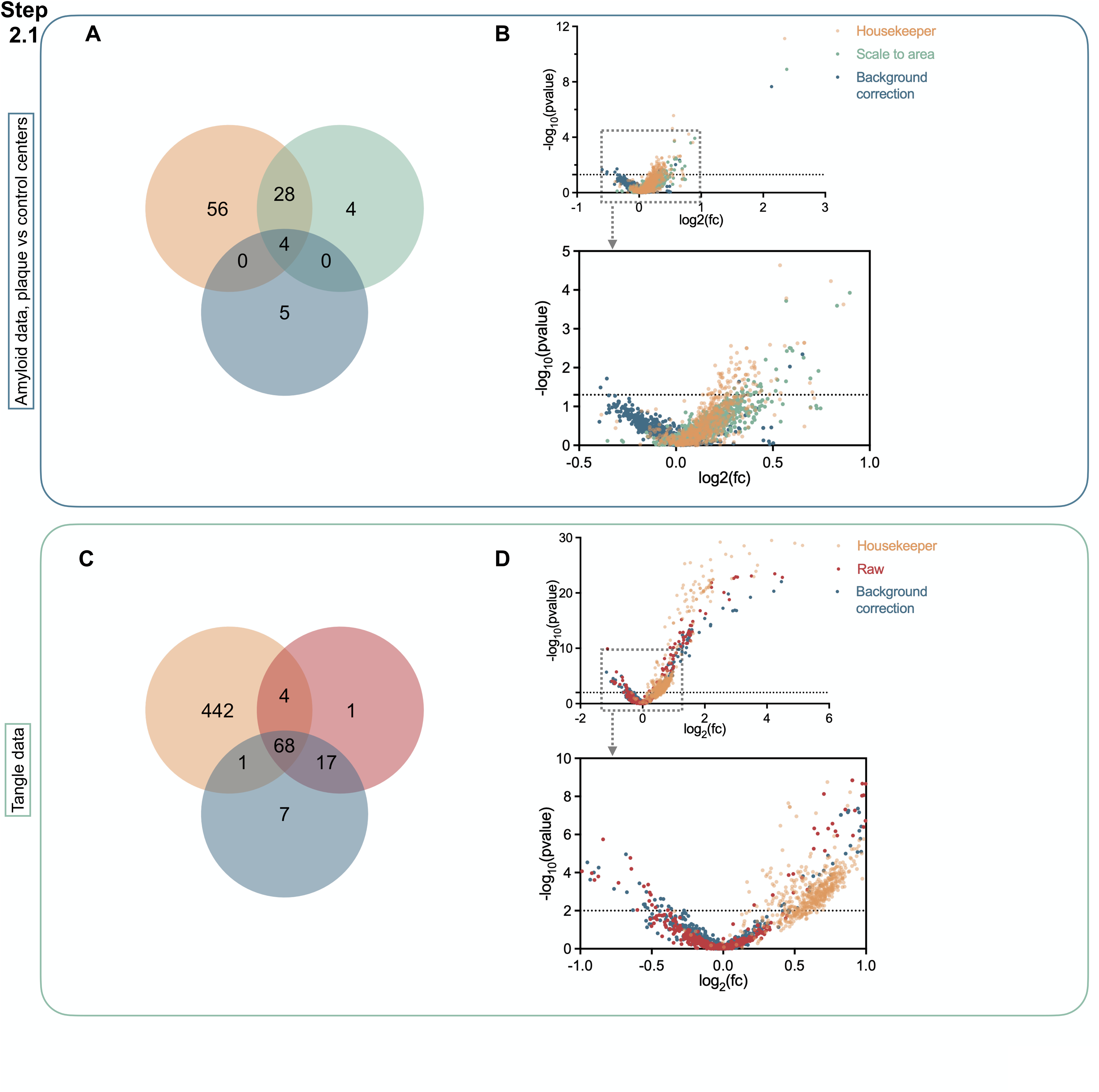
Impact of normalization strategies on downstream analysis. A. Overlap of DEPs identified in Center AOIs under three normalization strategies. For the plaque dataset, protein expression was normalized using one of three methods: housekeeping normalization (yellow), scaling to area (green), and background correction using IgGs (blue). Differential expression analysis compared Center AOIs from 6E10-positive regions to those from 6E10-negative regions. B. Overlay of volcano plots of DE analysis under three normalization strategies. DE analysis was performed on center AOIs. Statistical significance and effect size (fold change) varied depending on normalization strategies. C. Overlap of DEPs identified in the NFT single-neuron dataset under three normalization strategies. Protein expression was normalized using either housekeeping normalization (yellow), background correction with IgGs (blue), or analyzed as raw data (red). Differential expression analysis compared single neurons with and without NFT. D. Overlay of volcano plots of DE analysis under three data processing strategies. DE analysis was performed on NFT single-neuron dataset. Statistical significance and effect size (fold change) varied depending on normalization strategies.

### Biology-informed segment QC

Non-specific staining of morphology antibodies can lead to inclusion of ROIs that do not reflect relevant pathology. To ensure that selected AOIs captured authentic amyloid plaque features, we applied additional “biology-informed” segment QC following conventional baseline QC. We first quantified plaque load across all GeoMx-analyzed tissues. An auto-thresholding algorithm with size-exclusion criteria was applied to the 6E10 channel within three 2 mm × 2 mm regions per brain section. Non-specific signals were identified and manually excluded from final counts. Consistent with neuropathological diagnoses, no plaques were detected in NDC cases, whereas plaque counts in other subjects correlated with Thal staging (Fig. 4A). While 6E10 recognizes the C-terminal region of Aβ (Fig. 4B), bright, speckled signals in NDC tissue did not resemble true plaque morphology (Fig. 4C) and were excluded by automated thresholding (Fig. 4A).

**Figure 4.**
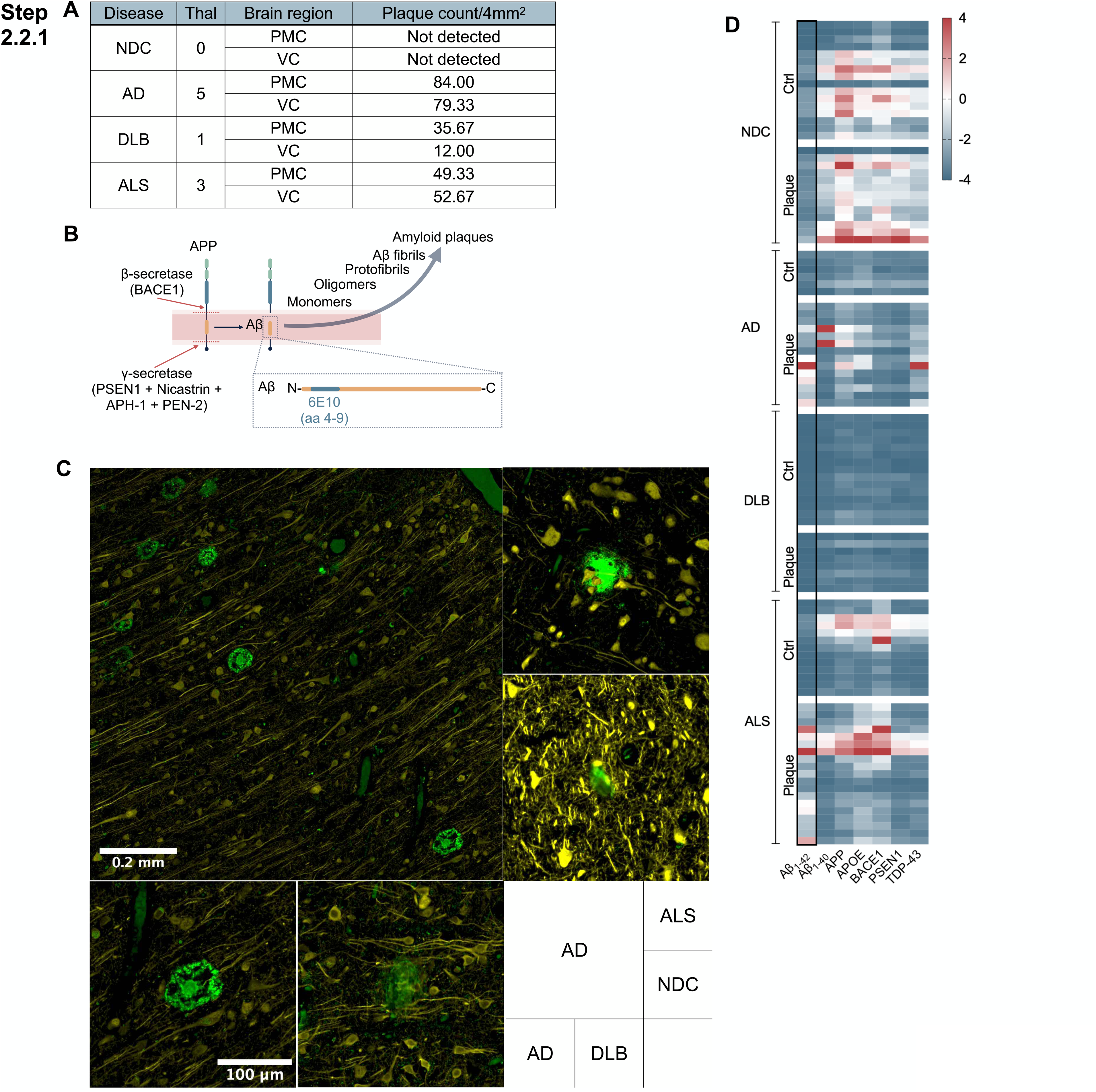
Biology informed segment QC: validation of plaque ROI selection. A. Quantification of amyloid plaque load across all tissue sections analyzed on the GeoMx DSP platform. Three 2 mm × 2 mm ROIs were drawn per section on 6E10-stained slides, and the average number of semi-automatically detected plaques was reported. B. Schematic illustrating APP processing and Aβ generation, including the epitope recognized by the 6E10 antibody. C. Representative images of 6E10 (green) staining from each case, highlighting differences in plaque morphology and distribution. Hud/MAP2 counterstaining is in yellow, D. Heatmap of selected Gx panel proteins associated with amyloid pathology. Expression values were scaled to area, and AOIs with extreme outlier values were temporarily excluded during scaling, then reintroduced and capped at a value of 4 for visualization (see Methods for details).

For the remaining cases, we validated that 6E10-positive regions corresponded to true plaques by examining a subset of proteins from the G_x_ panel, including Aβ_1-42_ and known plaque co-aggregate TDP43. Normalized counts confirmed that AOIs from AD and ALS cases exhibited molecular features consistent with true plaques (Fig. 4D). Independent staining of serial ALS sections using the orthogonal CellScape platform further confirmed plaque presence (Fig. 5). Based on these evaluations, all 6E10-positive AOIs from AD and ALS cases were retained for downstream analyses, whereas AOIs from NDC and DLB cases were excluded. These biology-informed QC steps ensured that subsequent analyses focused on ROIs accurately representing amyloid plaque pathology.

**Figure 5.**
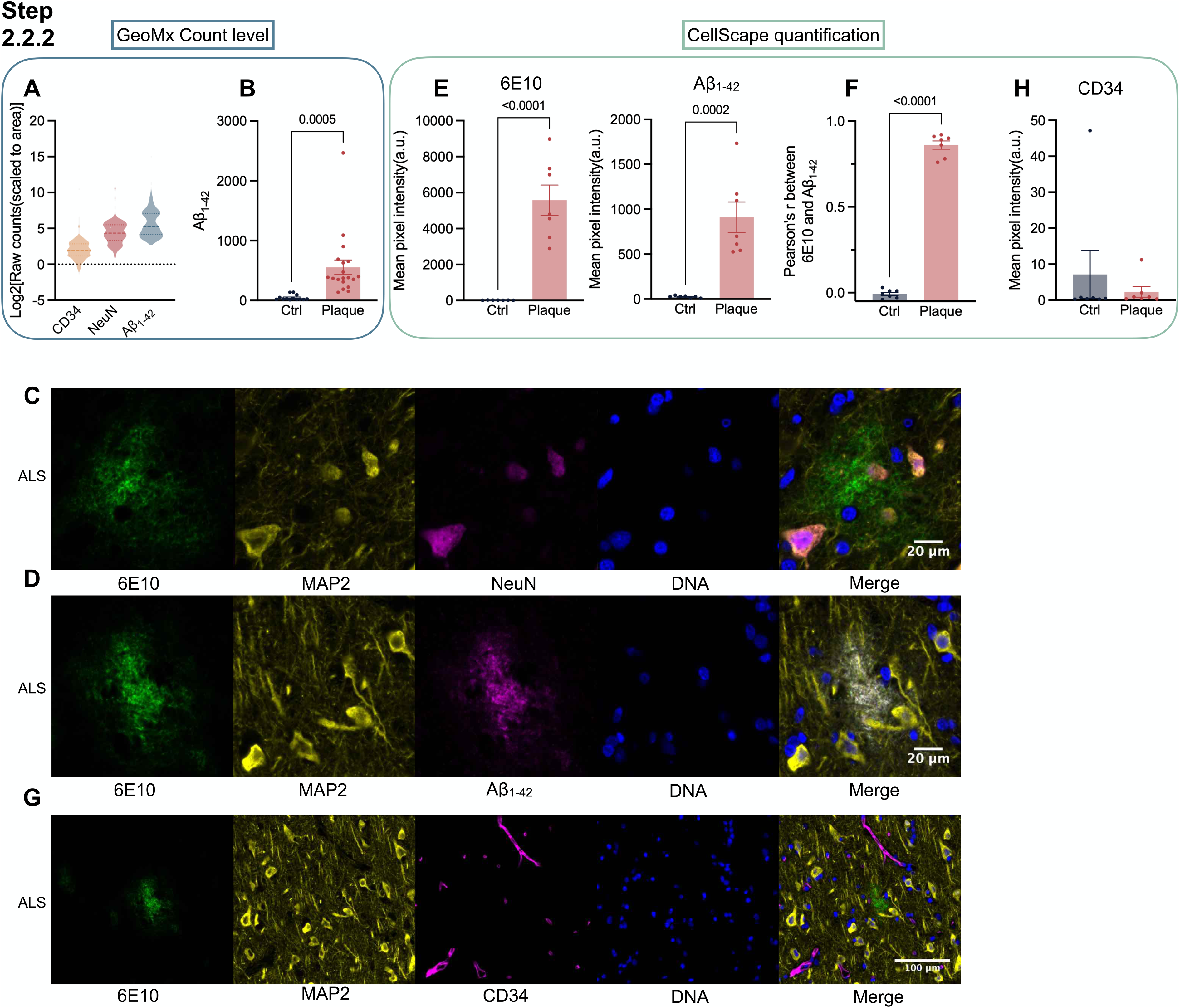
Biology informed target QC: Validation of selected G_x_ targets on CellScape. A. Signal strength of selected G_x_ targets from the GeoMx protein readout. Probe counts were scaled to the surface area of each AOI and then log2 transformed B. Aβ_1-42_ expression level measured by GeoMx. Each dot represents an AOI from either control or plaque center in the ALS case. Significance was determined using unpaired two-tailed t-test; error bar represents SEM. C. Representative CellScape staining of a serial section from the ALS PMC. Plaques were labeled using the 6E10 antibody (green). Neurons were labeled with MAP2 (yellow) and NeuN (magenta). D. Representative CellScape staining of a serial section from the ALS PMC. Plaques were labeled using the 6E10 antibody (green) and an antibody against Aβ_1-42_ (magenta). Neurons were labeled with MAP2 (yellow). E. Quantification of 6E10 and β-Amyloid_1-42_ staining. Plaque region was outlined by automated thresholding algorithm in Fiji. Control regions were selected by drawing circles of comparable size in plaque-free areas. Mean pixel intensity was measured for each region. Significance was determined using unpaired two-tailed t-test; error bar represents SEM. F. Colocalization of 6E10 and β-Amyloid_1-42_ staining. Pearson’s correlation coefficient (r) was calculated to describe the spatial colocalization between the two markers. Colocalization analysis was performed using Fiji’s Coloc2 plugin. G. Representative CellScape staining of CD34 (magenta) on the serial section from the ALS PMC. Plaques were labeled using the 6E10 antibody (green). Neurons were labeled with MAP2 (yellow). H. Quantification of CD34 signal within the plaque and control region as described in (B).

### Biology-informed target QC and validation of selective targets from the G_x_ panel with CellScape

G_x_ protein targets exhibited a broad dynamic range on GeoMx DSP (Fig. S4). To validate selected targets across this range, we stained serial sections from the same FFPE blocks on CellScape. Antibodies included MAP2, NeuN, 6E10, CD34, and Aβ1-42, using the same clones as in the G_x_ panel. MAP2, NeuN, and 6E10 were used for morphology, while CD34, NeuN, and Aβ_1-42_ represented low and high expressers on GeoMx, ensuring validation across differing signal intensities (Fig. 5A).

Neuron staining with MAP2 and NeuN was robust on CellScape, with morphology comparable to GeoMx readouts (Figs. 5A, C). Aβ_1-42_ signals co-localized with 6E10 in the ALS case, consistent with GeoMx measurements (Figs. 5C–E). CD34, a marker of vascular endothelial cells^21^, exhibited generally low counts in GeoMx AOIs, which could reflect either low antibody sensitivity or sparse vasculature (Figs. S4, 5A). On CellScape, CD34 staining clearly delineates the vasculature (Figs. 5G, H). This comparison indicates that low GeoMx CD34 counts correspond to regions with sparse CD34 cells, whereas high counts reflect dense vascularization.

Based on these validations, we retained all G_x_ protein targets for downstream analysis rather than pruning low-count targets. This approach preserves biologically meaningful variation and ensures that both low- and high-abundance proteins inform spatially resolved tissue features.

### Proteomic data renormalization and transcriptomic data processing

Proteomic data can require renormalization depending on AOI selection and the normalization strategy employed. Following biology-informed segment and target QC, only AOIs representing non-specific 6E10 staining were excluded. Counts had already been scaled to the minimum AOI surface area (from an AD case), and no further renormalization was performed. This approach preserved biologically meaningful variation while avoiding the introduction of normalization artifacts that could distort downstream analyses.

### Biological insights: RNA and protein concordance

With AOIs confirmed to reflect authentic plaque-associated regions, we processed WTA data on the Rosalind.bio platform. Center AOIs were excluded due to low cell and transcript counts, leaving 79 Ring1 and 83 Ring2 AOIs for downstream analysis. Because RNA and protein were measured from the same ROIs, we directly compared abundance for QC-passed segments. Antibodies recognizing post-translational modifications or targeting multiple proteins (e.g., pan-lamin A/B/C) were excluded, resulting in 223 RNA/protein pairs. Pearson correlations across all AOIs revealed positive, negative, and near-zero correlations (Figs. 6A–B). Positively correlated pairs included *MAP2*, *MBP*, *UCHL1*, and *BACE1*, reflecting essential neuronal functions and neuropathology. In contrast, *RPS6, FN1*, *MME*, and *FAS* were among the most negatively correlated pairs (Fig. 6C), highlighting cases where protein abundance was largely decoupled from transcript levels or subject to opposing regulatory mechanisms.

**Figure 6.**
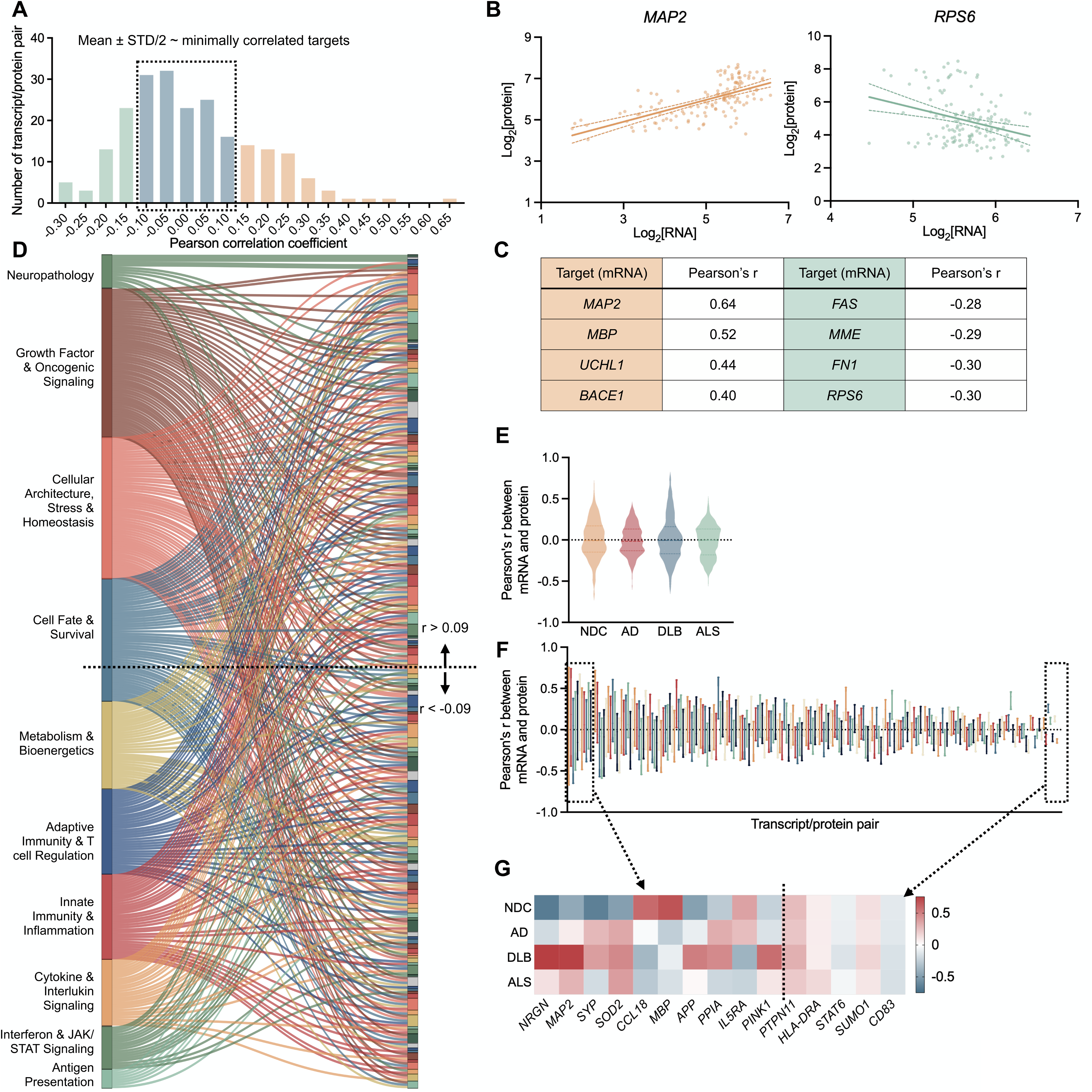
Concordance between protein and transcript readouts in GeoMx DSP. A. Histogram showing the distribution of Pearson’s r values for all matched RNA–protein pairs. All AOIs passed biology-informed QC in both RNA and protein analysis workflow were included in the analysis. B. Scatter plots showing the concordance between protein and transcript of *MAP2*(left) and *FN1*(right), respectively. Normalized RNA and protein counts were log_2_-transformed prior to plotting. The line represents a linear interpolation fit. C. Pearson’s r values for the top five most positively and most negatively correlated RNA–protein pairs. D. Sankey plot summarizing functional categories of RNA–protein pairs with some correlation strength. Bruker annotations were consolidated into ten functional groups (left), with targets sorted by Pearson’s r values (right). Targets with more than one functional annotations were shown in multiple links. E. Violin plot showing the distribution of Pearson’s r values for all matched RNA–protein pairs. AOIs passed biology-informed quality control in both RNA and protein workflows were stratified by disease condition. F. Box-and-whisker plots showing the range (min to max) of Pearson’s r values for each RNA–protein pair across disease conditions. Each line represents one matched pair. G. Heatmap illustrating protein/transcript pairs with the highest and lowest variance in Pearson’s r values across disease conditions.

Pairs with some correlation strength (defined as above or below the mean ± 0.5 SD, chosen to capture intermediate relationships) fell within a wide range of functional categories, including neuropathology, metabolism and bioenergetics, and cell fate and survival. However, no clear trend emerged linking strongly positive or negative correlations to any specific functional group (Fig. 6D). Minimally correlated pairs (within mean ± 0.5 SD) spanned diverse functional categories as well. Genes that encode key neurodegeneration-associated proteins such as *APP, MAPT,* and *TARDBP* exhibited complete decoupling between transcript and protein abundance. Important signaling molecules, such as *TP53*, *STING1*, *BCL2*, and *H2AX*, also showed minimal RNA–protein correlation (Fig. S5).

Stratification by disease condition revealed similar overall distributions of correlation coefficients (Fig. 6E), but individual targets often displayed marked shifts across conditions, indicating that RNA/protein relationships may be disease dependent (Fig. 6F). For instance, *NRGN* (neurogranin) exhibited strong RNA–protein correlation specifically in the DLB case but showed minimal or negative correlation across other disease conditions. Conversely, *SOD2* displayed strong positive correlation in the diseased individuals while displaying negative correlation in the NDC case (Fig. 6G).

### Biological insights: G_x_ and WTA panels detected disease-state dependent changes

We next evaluated whether this multi-omic approach captures molecular responses associated with plaque pathology. DE analysis of G_x_ proteomics data, using a linear mixed model, compared AOIs from plaque cores to matched control regions. As expected, Aβ_1-42_ was strongly upregulated in plaque center AOIs, confirming that the assay accurately detected the core pathological signature (Fig. 7A). Other DEPs included proteins involved in inflammation, antigen presentation, and glial activation (Fig. 7A). In contrast, the surrounding microenvironment exhibited relatively few changes (Fig. 7B), suggesting a localized molecular signature in which immune cells are tightly associated with plaque cores. Comparing DE results from Center AOIs before and after biology-informed QC revealed that this additional step not only changed the number and identity of DEPs but also altered their statistical significance and fold-change magnitudes, underscoring the importance of biology-informed ROI selection (Fig. S6).

**Figure 7.**
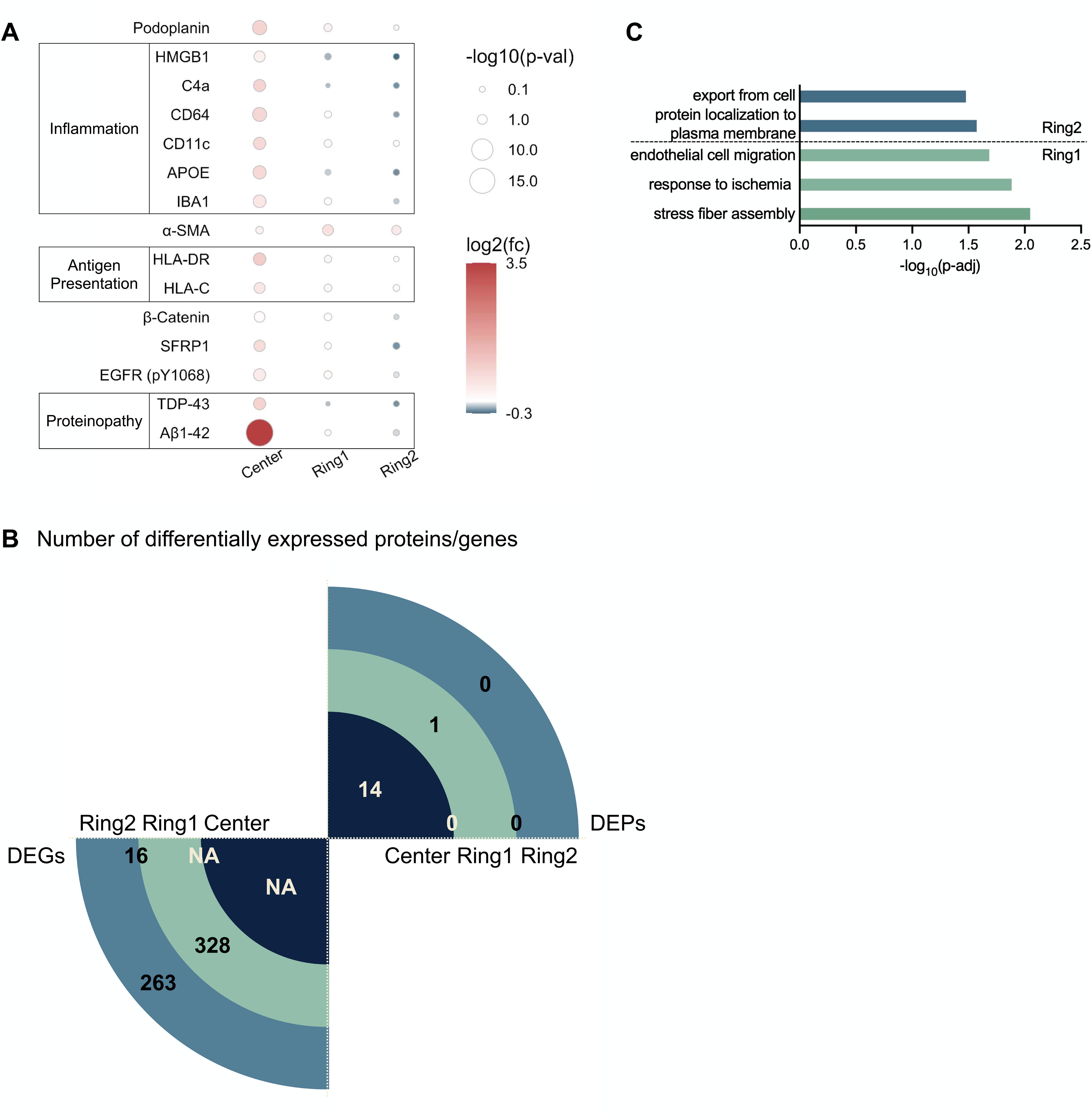
GeoMx multi-omics profiling revealed disease-state dependent molecular changes. A. DEPs between plaque and control AOIs. AOIs from all disease conditions were included in the analysis. Dot size represents the –log_10_(p-value), and color intensity corresponds to the log_2_(fold change). Differentially expressed proteins (p < 0.05) identified in each spatial segment were also displayed in the other two segments. Probe counts were scaled to the surface area of each AOI. B. Number of DEPs and DEGs in Center, Ring1, and Ring2 AOIs. For DEPs, statistical analysis was performed using the GeoMx Analysis Suite, including AOIs from all disease conditions. DEPs were defined by a p-value < 0.05. No DEPs were detected in Ring2 AOIs. For DEGs, statistical analysis was performed using the Rosalind.bio platform, including AOIs from all disease conditions. DEGs were defined by a p-value < 0.05 and an absolute fold change > 1.25. C. Functional enrichment analysis of top DEGs in the immediate environment surrounding plaques. DEGs were ranked by fold change, and analysis was performed using g:Profiler with default settings. Pathway enrichment was conducted within Gene Ontology: Biological Process. Adjusted p-values were corrected for multiple testing using the g:SCS algorithm. Selective pathways were shown.

DE analysis of WTA data on the Rosalind.bio platform revealed only modest overlap in differentially expressed genes (DEGs) between Ring1 and Ring2 AOIs (Fig. 7B). Gene ontology analysis using g:Profiler indicated that Ring1 DEGs were enriched for pathways involved in endothelial cell migration, consistent with the observed upregulation of α-SMA protein, a marker of pericytes and endothelial cells within the plaque core, as well as response to ischemia and stress fiber assembly. In contrast, Ring2 DEGs were enriched for pathways related to protein export and plasma membrane localization, reflecting spatially distinct alterations in secretion and membrane-associated signaling (Fig. 7C). Together, these findings demonstrate that integrated proteomic and transcriptomic profiling captures a spatially resolved gradient of molecular responses to plaque pathology, providing a multi-modal view of localized immune, vascular, and cellular signaling changes in neurodegenerative disease.

## Discussion

This study establishes a biology-informed framework for applying spatial multi-omics to postmortem brain tissue across neurodegenerative diseases. By integrating high-plex proteomic and transcriptomic profiling with rigorous normalization, quality control, and orthogonal validation, we address a central limitation in current spatial biology workflows — the lack of standardized, reproducible analytical approaches. This framework preserves biologically meaningful variation, enables accurate RNA–protein integration, and supports robust interpretation of complex tissue organization in both health and disease.

Protein normalization is a critical first step. Protein signals are sensitive to AOI/ROI-specific factors, antibody performance, and background noise, where errors at this stage can propagate into downstream RNA–protein integration. We evaluated commonly used normalization strategies, including housekeeping proteins, IgG negative controls, and AOI/ROI area scaling. Housekeeping proteins exhibited disease-dependent variation in both plaques and tau tangles, underscoring that conventional internal references may not be stable in pathological contexts. IgG signals were inconsistent across AOIs/ROIs and did not reliably reflect non-specific background binding. For plaque AOIs, scaling to surface area produced the most consistent results, whereas single-neurons were best corrected using ROI-specific signal-to-background adjustment. Additionally, to enable cross-segment comparisons in downstream analyses, we normalized all plaque AOIs together. Alternatively, AOIs can be separated into three groups and normalized independently using signal-to-background correction, but only if cross-segment comparison is not part of the analysis design. These findings emphasize the importance of context-specific evaluation of protein normalization prior to RNA–protein integration, preserving biologically meaningful variation while minimizing artifacts.

Beyond normalization, biology-informed QC was essential for robust analysis. Conventional baseline QC alone can miss tissue artifacts, non-specific antibody binding, and high endogenous autofluorescence, all of which may obscure true pathological signals. By applying segment- and target-level QC guided by morphology, orthogonal validation (CellScape), and pathological context, we ensured that downstream analyses were performed on biologically meaningful, pathologically relevant features rather than technical or anatomical artifacts.

Integration of RNA and protein measurements revealed overall poor concordance. A subset of protein targets exhibited minimal or even negative correlation with their corresponding transcripts, and the strength of RNA–protein concordance varied by disease condition. This highlights the risk of inferring protein-level changes from transcriptomic data alone and reinforces the value of integrated multi-omic profiling for accurate biological interpretation. Protein data provided robust detection in plaque cores, where RNA was sparse, revealing glial and immune cell activation tightly associated with plaques. Transcriptomic profiling of surrounding microenvironments captured region-specific gene expression changes across diverse cell types, enabling complementary insights into spatially resolved molecular dynamics.

Importantly, this dataset itself represents a valuable resource for the community. It offers high-plex, spatially resolved RNA and protein measurements across multiple neuropathological features and brain regions, enabling further exploration of RNA–protein relationships, immune and vascular microenvironments, and disease-specific molecular signatures. These data can serve as a benchmark for validating analytical approaches and for future studies leveraging spatial proteomics and integrated multi-omics to interrogate complex tissues.

Despite these advances, limitations should be acknowledged. The study included a limited number of human cases and brain regions, and our observations should be validated in independent cohorts. Our focus was on currently available normalization and QC strategies, and new computational and experimental approaches will likely continue to emerge. The primary goal was to provide broadly applicable, biology-informed guidelines for rigorous data processing rather than to establish definitive biological conclusions.

Overall, this work demonstrates that rigorous, biology-informed QC, normalization, and validation of spatial proteomic and transcriptomic data are important for reliable interpretation of high-plex multi-omic assays. By establishing and validating these workflows, we provide a framework that can be broadly applied across studies, enabling reproducible and accurate exploration of complex tissue organization and molecular pathology in neurodegenerative disease and beyond.

## Method

### Human samples

Brain tissues from the cases profiled were provided by the University of Washington’s Biorepository and Integrated Neuropathology Laboratory. The neuropathologic diagnoses given at the time of original autopsy evaluation, hematoxylin and eosin (H&E), and immunohistochemical (IHC) stains were reviewed by board-certified neuropathologist C.D.K to ensure consistency in diagnoses.

### Tissue section preparation and ROI selection for GeoMx multi-omics profiling

FFPE sections covering either the primary motor or visual cortices were deparaffinized and processed according to the manufacturer’s protocol, GeoMx DSP Spatial Proteogenomics Assay manual (Bruker, MAN-10158-05-01). The WTA panel was used for transcript detection, and the protein panel combined the Immuno-Oncology Proteome Atlas (IPA) with 63 spike-in proteins. Because STING was represented in both the IPA and the custom spike-in’s, the probe from the IPA panel was excluded from all downstream analyses to prevent redundancy. In total, we profiled 637 biological targets with 5 negative probes (IgGs). Tissue sections were incubated with a panel of fluorescently labeled morphology markers to facilitate ROI selection: tau phosphorylated at Ser202/Thr205 was used as a surrogate marker of neurofibrillary tangles (AT8 antibody, conjugated to Alexa Fluor 594 (AF594)^22^; amyloid plaques were detected with the 6E10 antibody conjugated to AF488; neuronal markers HuD and MAP2 were co-labeled with AF647; nuclei were visualized using SYTO 83 nucleic acid stain. For amyloid plaques, a circular ROI was drawn around each plaque in cortical layer 5, with two concentric rings to capture the surrounding microenvironment. For tau tangles, a 25 μm-diameter circle centered on the cell body was drawn. The total number of ROIs/AOIs is reported in Figure 1.

Plaque ROI selection was verified by post-collection at the biology-informed QC step. Plaque load was quantified using Fiji. Multichannel whole tissue scans were exported from the GeoMx DSP platform, and plaque counts were performed on the 6E10 channel. For each sample, three ROIs (2 mm × 2 mm each) were analyzed per tissue section, and the mean value was reported as plaque count per 4 mm². An appropriate auto global threshold and size exclusion criteria were applied for each sample to isolate plaque-specific signals. Non-specific signals captured during this process were manually reviewed and excluded from the final plaque count.

### GeoMx multi-omics data processing

The G_x_ protein panel data were processed using the GeoMx Analysis Suite software, with quality control and normalization details provided in the Results section. For the conventional baseline QC, the following exclusions were applied: In the plaque dataset, one AOI was excluded due to low sequencing saturation, one AOI was excluded due to low surface area, and three AOIs were excluded due to uniformly low read counts (read value of 1 across all targets). In the single-neuron dataset, three ROIs were excluded due to low raw read counts, two ROIs were excluded due to a low percentage of aligned reads, and one ROI was excluded due to abnormally high raw read counts across all targets. For the biology-informed QC, all plaque centers and associated rings in DLB and NDC cases were excluded.

WTA data were processed using the GeoMx DCC (WTA) Methods workflow in Rosalind.bio following default quality control (QC) metrics. Because non-template controls (NTCs) were not included during sample collection, AOIs that failed sequencing were reassigned as NTCs to ensure compatibility with the Rosalind analysis pipeline. Only AOIs that passed biology-informed QC metrics were retained for downstream analysis. Grubbs’ outlier test was applied to identify and remove outlier negative probes. The Limit of Quantification (LOQ) was calculated per AOI as two standard deviations above the geometric mean of the negative probes. Gene detection threshold was set at 0% to select genes above the LOQ for AOI performance evaluation and all center AOIs regardless of their detection level were excluded at this step. An AOI detection rate threshold of 5% was applied to identify AOIs with sufficient gene detection for assessing gene performance. Quantile normalization was used to normalize gene expression values across AOIs.

Heatmaps were generated using area-scaled data normalized to a range of -4 to 4 for visualization per ROI or AOI. Because background intensity was measured across the entire ROI, IgG and negative WTA probe counts were calculated as the geometric mean of the three AOIs within each ROI, representing the ROI-wide background values (Fig. 2F). Control and plaque center AOIs were used to generate a heatmap of plaque-associated proteins with outlier AOIs temporarily excluded during -4 to 4 scaling and reintroduced as a capped value of 4 for plotting (Fig. 4D). Specifically, ALS PMC (Aβ1-42 and APOE), AD VC (Aβ1-40), and NDC PMC (APP) plaque center AOIs were initially excluded to improve visualization of relative protein expression across AOIs.

Concordance analysis was performed on the plaque dataset using AOIs that passed biology-informed QC in both RNA and protein workflows. Antibodies recognizing post-translational modifications or multiple proteins were excluded. Protein targets were matched to WTA transcripts by HUGO gene symbol, and Pearson’s *r* values were computed using the R “cor” function.

### Differential expression analysis

DE analysis of the G_x_ data was performed using the GeoMx Data Analysis Suite software. Protein expression between plaque-positive and plaque-negative AOIs (Center, Ring1, and Ring2, respectively) was compared using a linear mixed model (LMM), followed by Benjamini-Hochberg correction to determine log_2_ fold changes and associated p-values. A random intercept for Scan ID was included in the model to account for variability across tissue sections. Differentially expressed proteins with p-value < 0.05 were reported.

DE analysis of the WTA data was performed using Rosalind.bio. A LMM was applied to estimate differential gene expression, log fold changes, and associated p-values. The model incorporated a random intercept for the assigned Tissue ID to account for variability between tissue samples. To identify significantly differentially expressed genes, we applied the following thresholds: absolute fold change > 1.25 and p-value < 0.05.

### Pathway analysis

Pathway enrichment analysis was done using the gprofiler2 package in R. Genes with a p-value < 0.05 from the differential expression analysis were included, and log fold change values were used to rank the genes. Gene set enrichment was performed using the Gene Ontology Biological Processes (GO: BP) database, with the default data source provided by g:Profiler. Multiple testing correction was applied using the default g:SCS method. Enriched GO: BP terms were visualized as bar plots, displaying the -log (p-value) of each term.

### Tissue section preparation and image analysis for CellScape staining

Serial sections covering the primary motor cortices from the same human subject were deparaffinized and processed according to the manufacturer’s protocol, CellScape Sample Preparation and Instrument Operation manual (Bruker, MAN-10200-02). Tissue sections were incubated with a panel of fluorescently labeled antibodies, using the same clones as those in GeoMx profiling: MAP2, NeuN, CD34, 6E10, Aβ_1-42_, and phosphorylated Tau (pTau S404).

ROIs centered on amyloid plaques and control regions, identified by 6E10 staining, were annotated using QuPath (v0.6.0) and exported as multi-channel TIFF images for further analysis in Fiji. Plaque masks were generated using the Moments auto-thresholding method, while control region masks were defined as circular areas of comparable diameter. Mean pixel intensities of 6E10, Aβ_1-42_, and CD34 were measured. The Pearson correlation coefficient between 6E10 and Aβ_1-42_ signals was calculated using the Coloc2 plugin in Fiji.

### Statistical analysis

Except DE and concordance analyses, all other statistical analyses were performed using Prism 10 software (GraphPad). Specific tests used for each figure are indicated in the corresponding figure legends.

## Data availability

The pre-quality control and supporting data for all experiments are available from the authors on reasonable request.

## Author contributions

Conceptualization: M.E.O. and T.C.O.; data acquisition: T.C.O., A.R., M.I., and O.B.; data analysis and interpretation: X.S., H.R.H., S.K., T.C.O, and M.E.O; writing—original draft: X.S.; editing and final review of the manuscript: all authors; technical and material support: C.D.K.; supervision: J.M.B. and M.E.O.; funding acquisition: M.E.O.

## Declaration of interests

M.E.O. has a patent pending, ‘Detecting and Treating Conditions Associated with Neuronal Senescence’ unrelated to this work. M.E.O. is the Director of a Bruker Spatial Biology Center of Excellence.

A.R., M.I., O.B., and J.M.B. are or were employees of Bruker Spatial Biology, Inc.

The other authors declare no competing interests in relation to this work.

## Acknowledgements

M.E.O. is supported by the Alzheimer’s Drug Discovery Foundation (GC-201908-2019443), Cure Alzheimer’s Fund, Hevolution/American Federation for Aging Research, National Institute on Aging (R01AG068293, R01AG085182, R21AG087907, R01AG0909551, R25AG073119; U54AG079754, R01AG079224), National Institute of Neurological Disorders and Stroke (R56NS131387), the Rainwater Charitable Foundation, and US Department of Veterans Affairs (I01BX005717).

The authors used ChatGPT (OpenAI, San Francisco, CA; GPT-5) for language editing and stylistic refinement. The authors take full responsibility for the content of the manuscript.

## Figure Legends

**Supplementary table 1 (related to figure 1). Information on human samples**

**Supplementary figure 1 (related to figure 1). Detailed QC metrics and data processing**

A. Raw counts from G_x_ analysis. Each dot represents an AOI.

B. Raw counts from WTA analysis. Each dot represents an AOI.

C. Number of detected genes from WTA analysis. Each dot represents an AOI.

D. Number of cells from each ROI. Syto83 nuclear staining was used for quantification. Significance was determined using Brown-Forsythe and Welch Anova tests with Dunnett T3 correction; error bar represents SEM.

E. Number of neurons from each ROI. MAP2+HuD staining was used for quantification. Significance was determined using Brown-Forsythe and Welch Anova tests with Dunnett T3 correction; error bar represents SEM.

**Supplementary figure 2 (related to figure 2). Results from the “Evaluate-Normalization-Options” DSP DA Script.**

A. Overall agreement of the housekeeping proteins. Each dot represents an AOI and colored by disease condition and brain region. The numbers on the plot refer to the correlation variability.

B. Overall agreement of IgGs. Each dot represents an AOI and colored by disease condition and brain region. The numbers on the plot refer to the correlation variability.

**Supplementary figure 3 (related to figure 2). Considerations of normalization for a single-cell dataset.**

G. Expression levels of housekeeping probes across distinct neuronal populations. Neurons harboring NFT and nearby tangle-negative neurons (Ctrl) were selected from cortical layers 2 (L2) and 5 (L5) of the ADwNP PMC sample.

H. Count levels of IgGs.

I. Count levels of IgGs across different neuronal populations.

J. Correlation between mouse IgG2b (used as a negative control probe) and other IgG probes. Each point represents an AOI. The zoomed-in panel (displaying AOIs with IgG counts below 25) highlights a loss of linearity and concordance at low count levels.

K. A representative image showing varying IgG levels in adjacent neurons. Neurons were stained with MAP2+HuD (yellow). Hyperphosphorylated tau species were stained with AT8 (magenta). Profiled neurons were highlighted by white circles and labeled with identifiers above. The corresponding raw IgG counts were provided in the table.

L. Correlation between IgG counts and total target counts for each ROI. The x-axis shows the geometric mean of IgG probe counts for each region of interest (ROI), while the y-axis represents the total raw counts of all protein probes in the corresponding ROI. Since the data was not normally distributed, Spearman’s correlation coefficient (R) and associated p-value are reported.

**Supplementary figure 4 (related to figure 5). Log□ signal-to-background ratio for all targets**

This plot was automatically generated using the “Evaluate-Normalization-Options” script in the DSP Data Analysis (DA) pipeline.

**Supplementary figure 5 (related to figure 6). Sankey plot summarizing functional categories of minimally correlated RNA-protein pairs**

Bruker annotations were consolidated into ten functional groups (left), with targets sorted by Pearson’s r values (right). Targets with multiple functional annotations were shown in multiple links; key targets are highlighted.

**Supplementary figure 6 (related to figure 7). The impact of biology informed QC on downstream analysis**

A. Overlap of DEPs identified in Center AOIs before and after biology-informed QC. Protein expression values were scaled to area prior to analysis. Differential expression analysis compared Center AOIs before (green) and after (yellow) exclusion of segments failing biology informed QC.

B. Overlay of volcano plots of DE analysis with Center AOIs before and after biology-informed QC. Statistical significance and effect size (fold change) varied depending on normalization strategies.

C. List of DEPs identified under both QC conditions. For each target, statistical significance and effect size (fold change) are reported.

