## Supplemental table and figures for "Spatial Multi-Omics Workflow and Analytical Guidelines for Alzheimer’s Neuropathology"

| Sample | Alzheimer's Disease (AD) | Dementia with Lewy Bodies (DLB) | Amyotrophic Lateral Sclerosis (ALS) | Non-demented Control (NDC) |
|---|---|---|---|---|
| Sex | M | M | F | F |
| Age at death | 80 | 73 | 72 | 56 |
| Post-mortem interval (hrs) | 8.3 | 7.4 | 9.8 | 6.3 |
| Cognitive status | Dementia | Dementia | No dementia | No dementia |
| Overall AD neuropathological change | High | Low | Low | None |
| Braak staging[1] | 6 | 1 | 1 | 1 |
| CERAD score[2] | 3 | 0 | 0 | 0 |
| Thal phase[3] | 5 | 1 | 3 | 0 |
| Overall CAA score | 0 | 0 | 0 | 0 |
| Highest Lewy Body disease | Not identified | Neocortical (Diffuse) | Not identified | Not identified |
| Primary neuropath diagnosis | AD | AD pathology present but insufficient for AD diagnosis; DLB, with or without AD | ALS, present with TDP-43 inclusions | Normal brain |

[1]Braak, Heiko, and Eva Braak. "Neuropathological stageing of Alzheimer-related changes." *Acta neuropathologica* 82.4 (1991): 239-259.
[2]Chandler, Melanie J., et al. "A total score for the CERAD neuropsychological battery." *Neurology* 65.1 (2005): 102-106.
[3]Thal, Dietmar R., et al. "Phases of Aβ-deposition in the human brain and its relevance for the development of AD." *Neurology* 58.12 (2002): 1791-1800.

Supplementary table 1

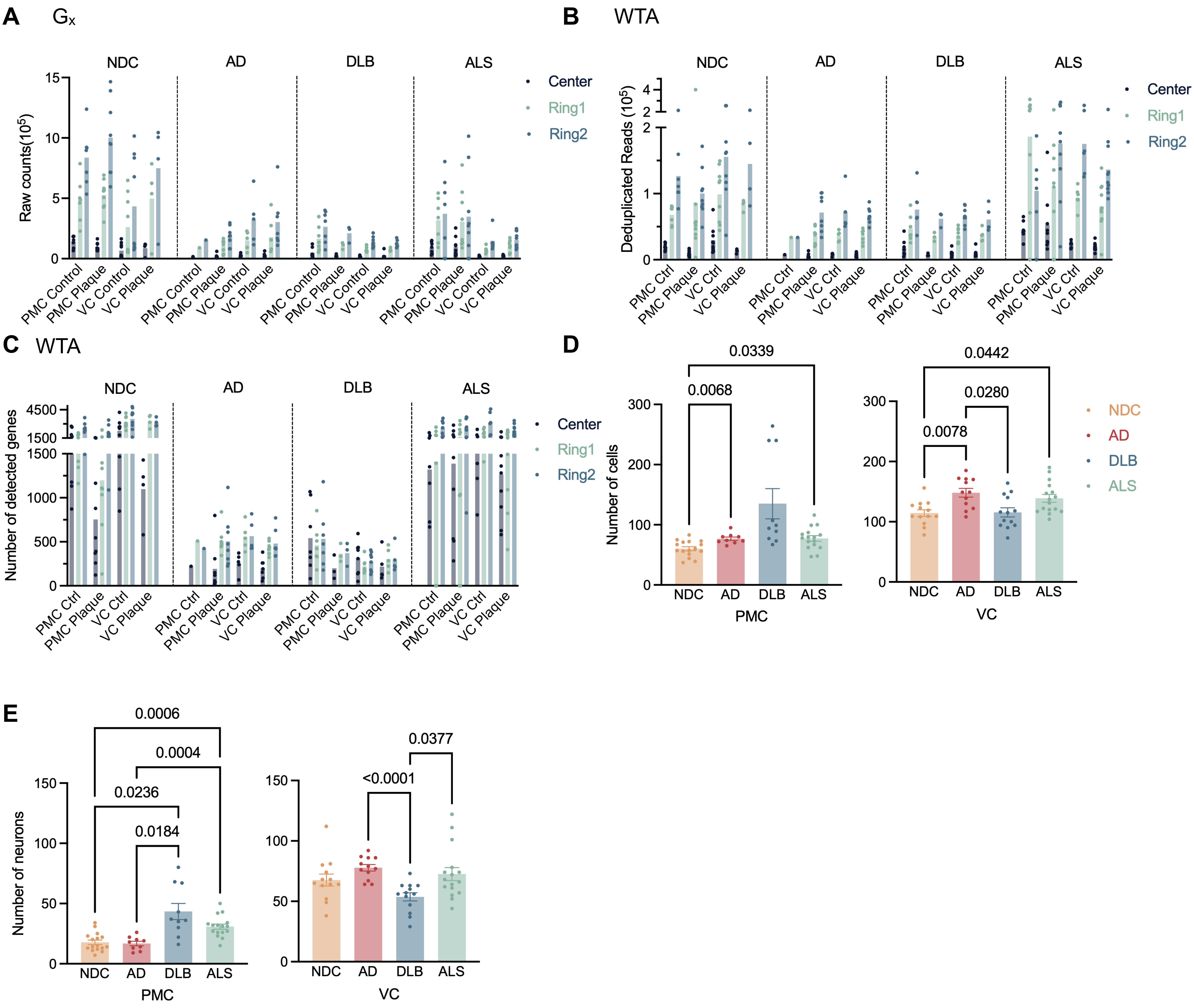


Supplementary Figure 1

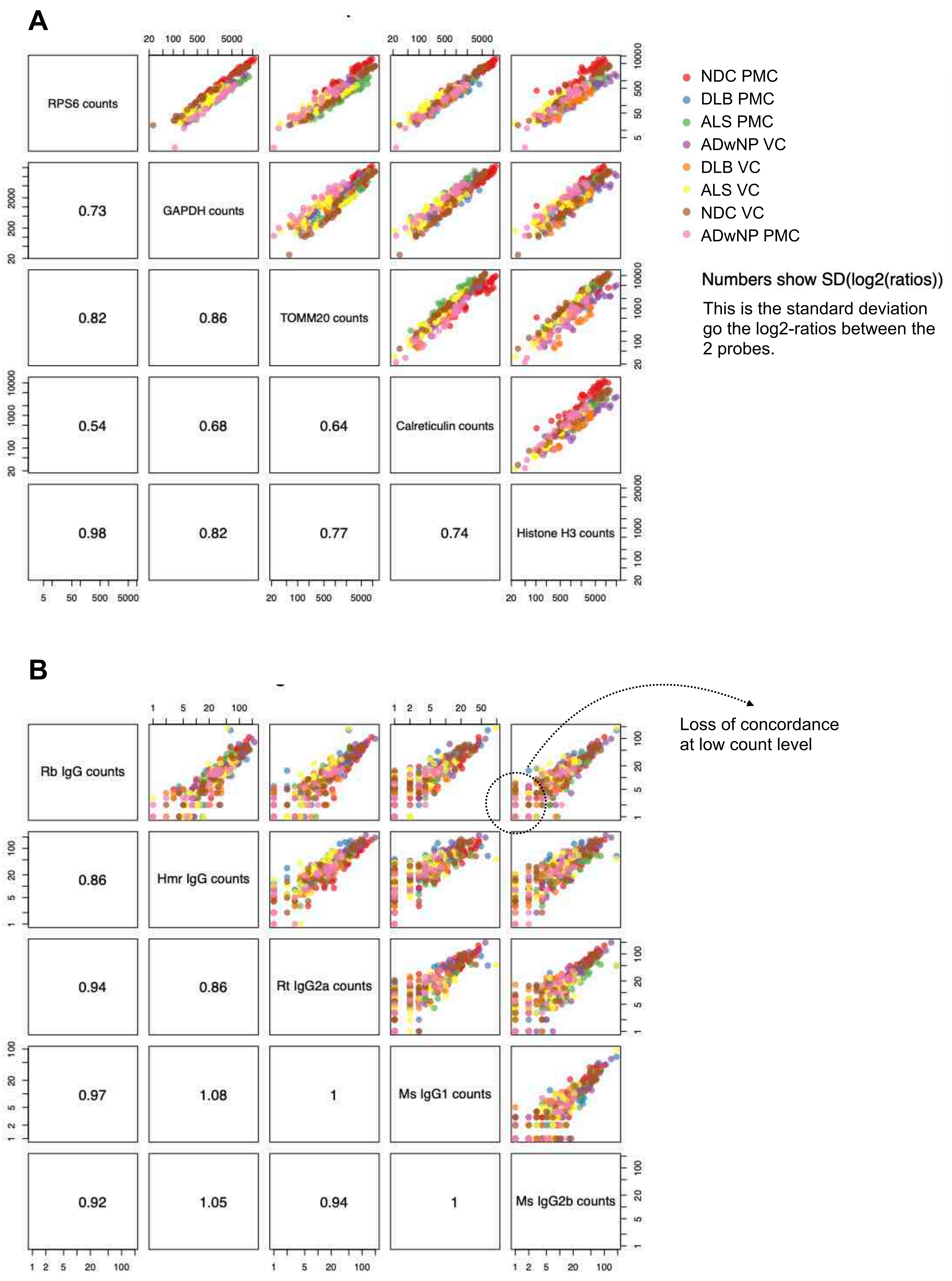
A
RPS6 counts
GAPDH counts
TOMM20 counts
Calreticulin counts
Histone H3 counts
0.73
0.82
0.86
0.54
0.68
0.64
0.98
0.82
0.77
0.74
NDC PMC
DLB PMC
ALS PMC
ADwNP VC
DLB VC
ALS VC
NDC VC
ADwNP PMC
Numbers show SD(log2(ratios))
This is the standard deviation go the log2-ratios between the 2 probes.
B
Rb IgG counts
Hmr IgG counts
Rt IgG2a counts
Ms IgG1 counts
Ms IgG2b counts
0.86
0.94
0.86
0.97
1.08
1
0.92
1.05
0.94
1
Loss of concordance at low count level

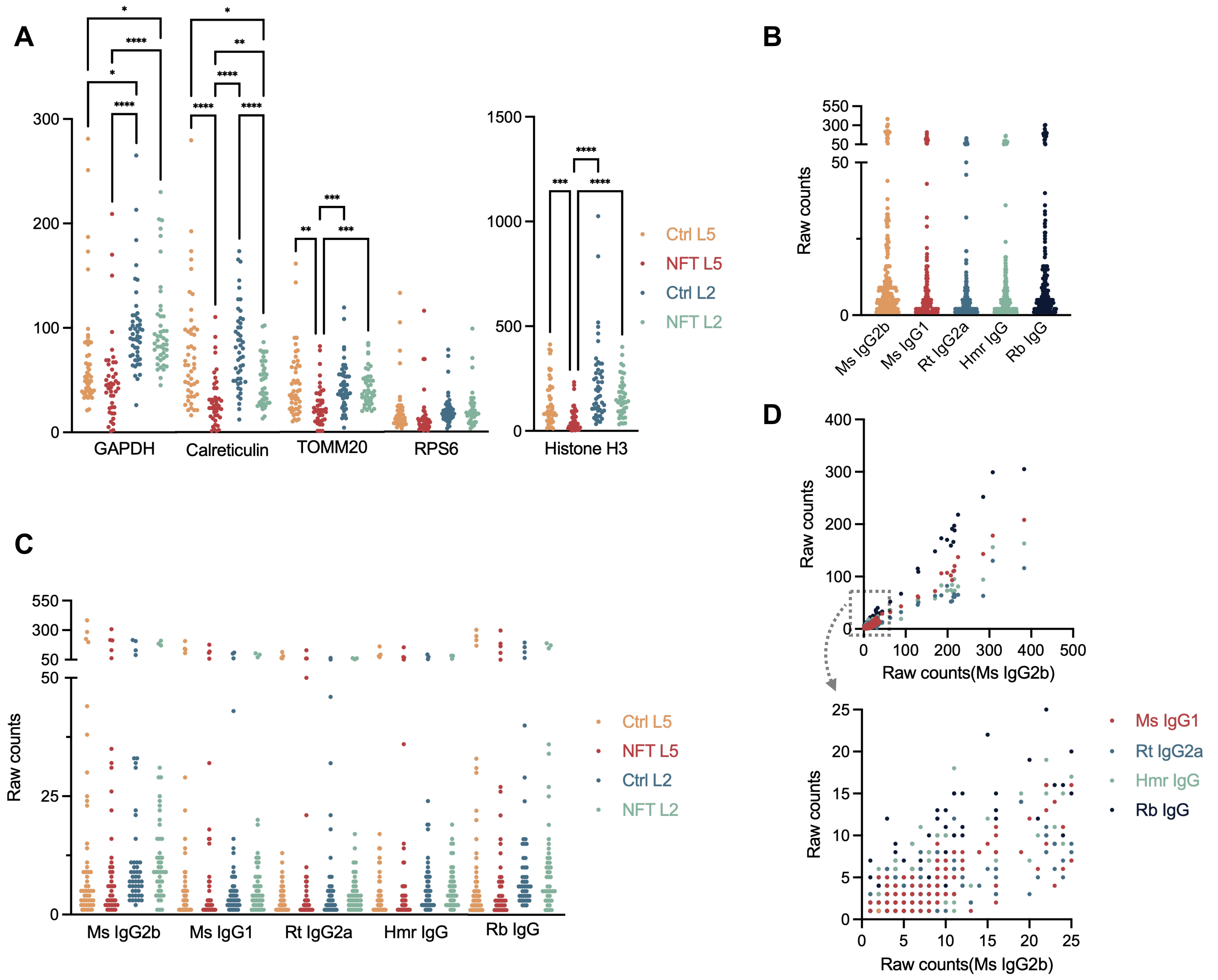


E

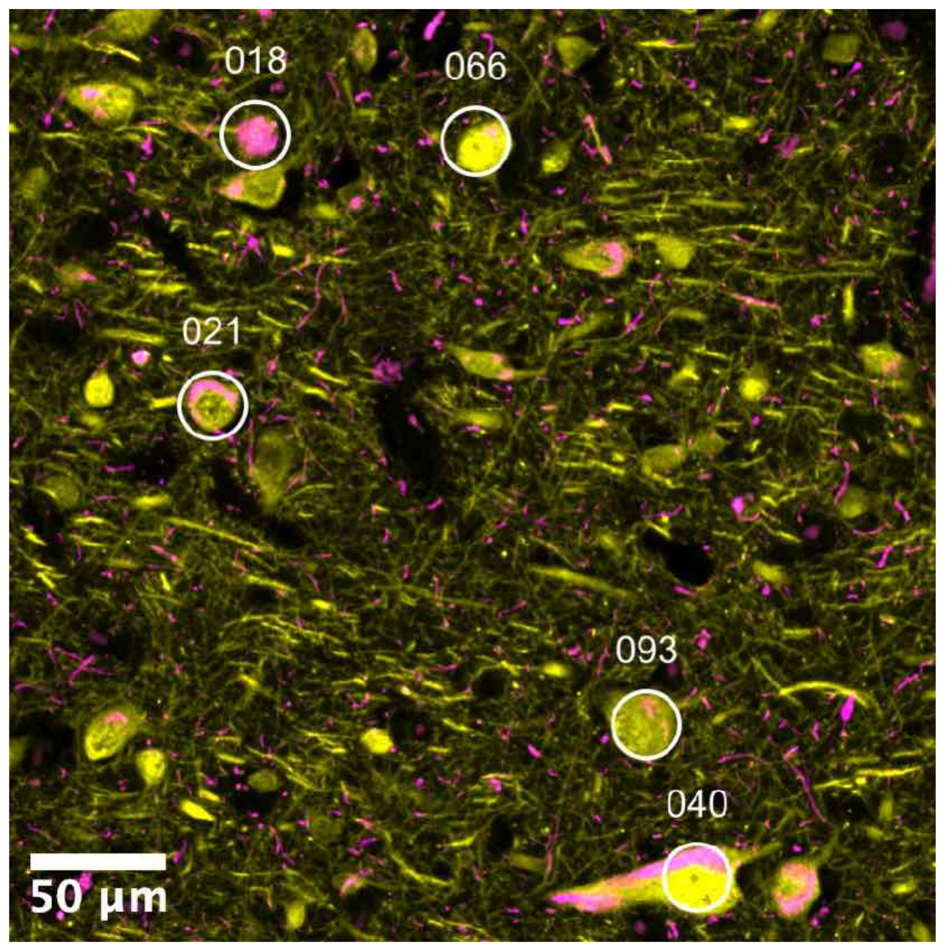


| ROIs | Ms IgG2b | Ms IgG1 | Rt IgG2a | Hmr IgG | Rb IgG |
|---|---|---|---|---|---|
| 18 | 308 | 178 | 130 | 156 | 299 |
| 21 | 2 | 1 | 1 | 1 | 2 |
| 40 | 3 | 1 | 1 | 1 | 4 |
| 66 | 285 | 143 | 63 | 94 | 252 |
| 93 | 5 | 5 | 2 | 4 | 4 |

F

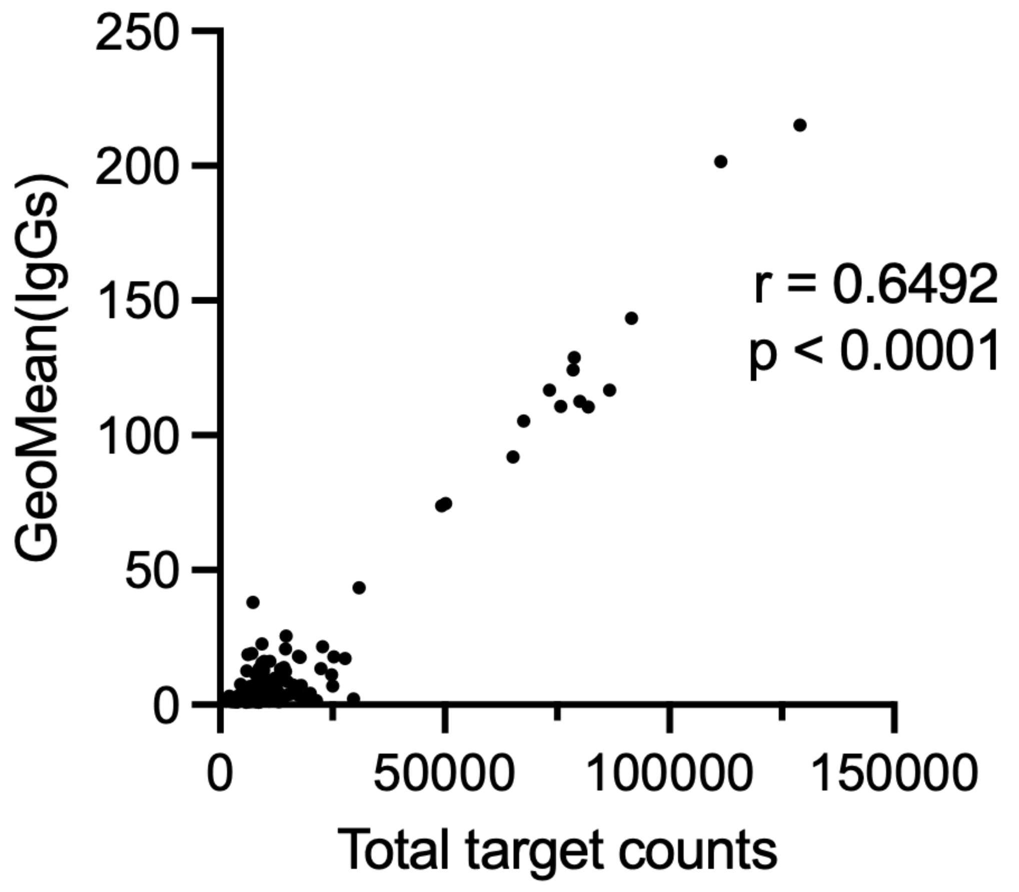


Supplementary Figure 3

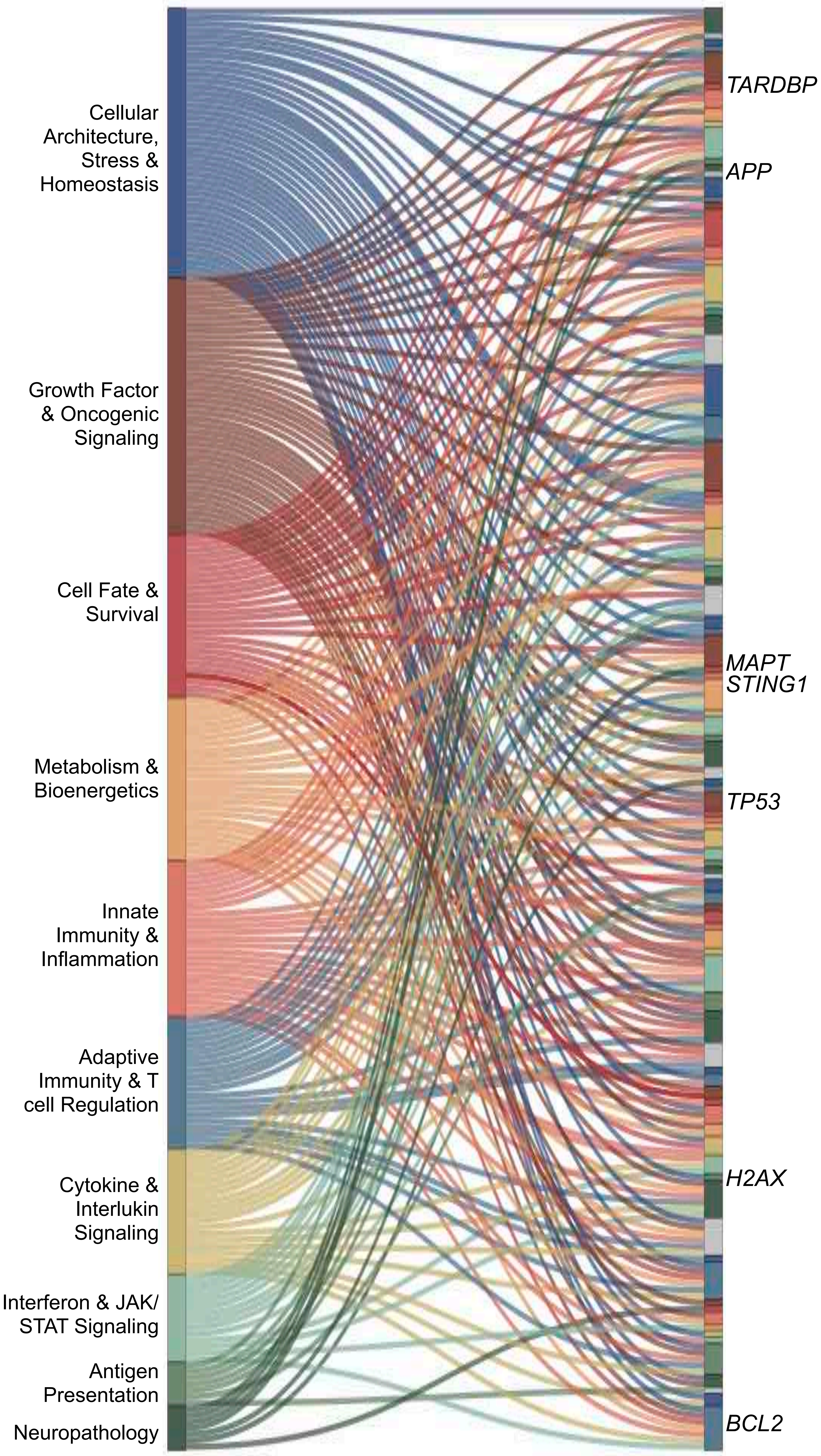


Supplementary Figure 5

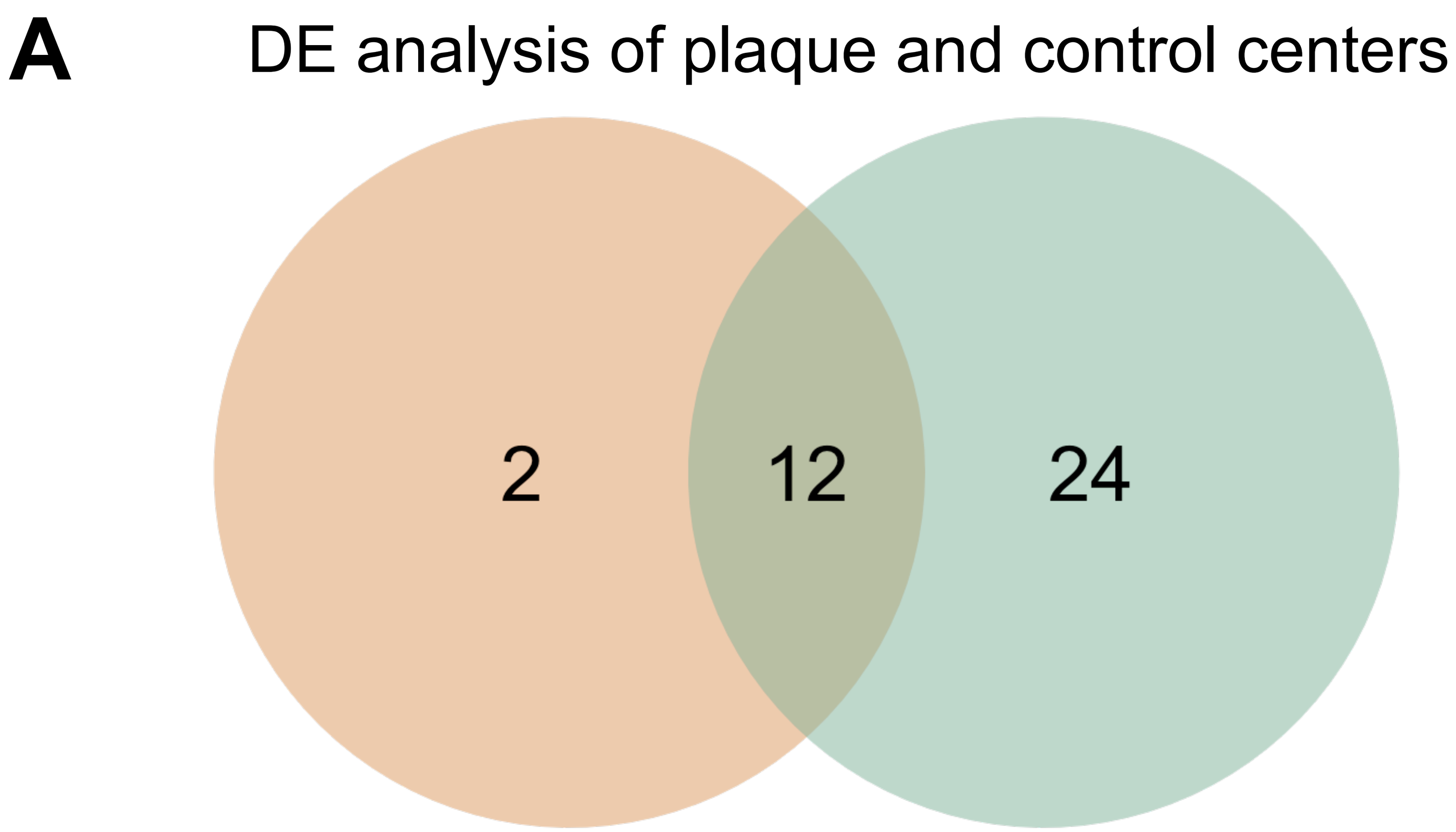


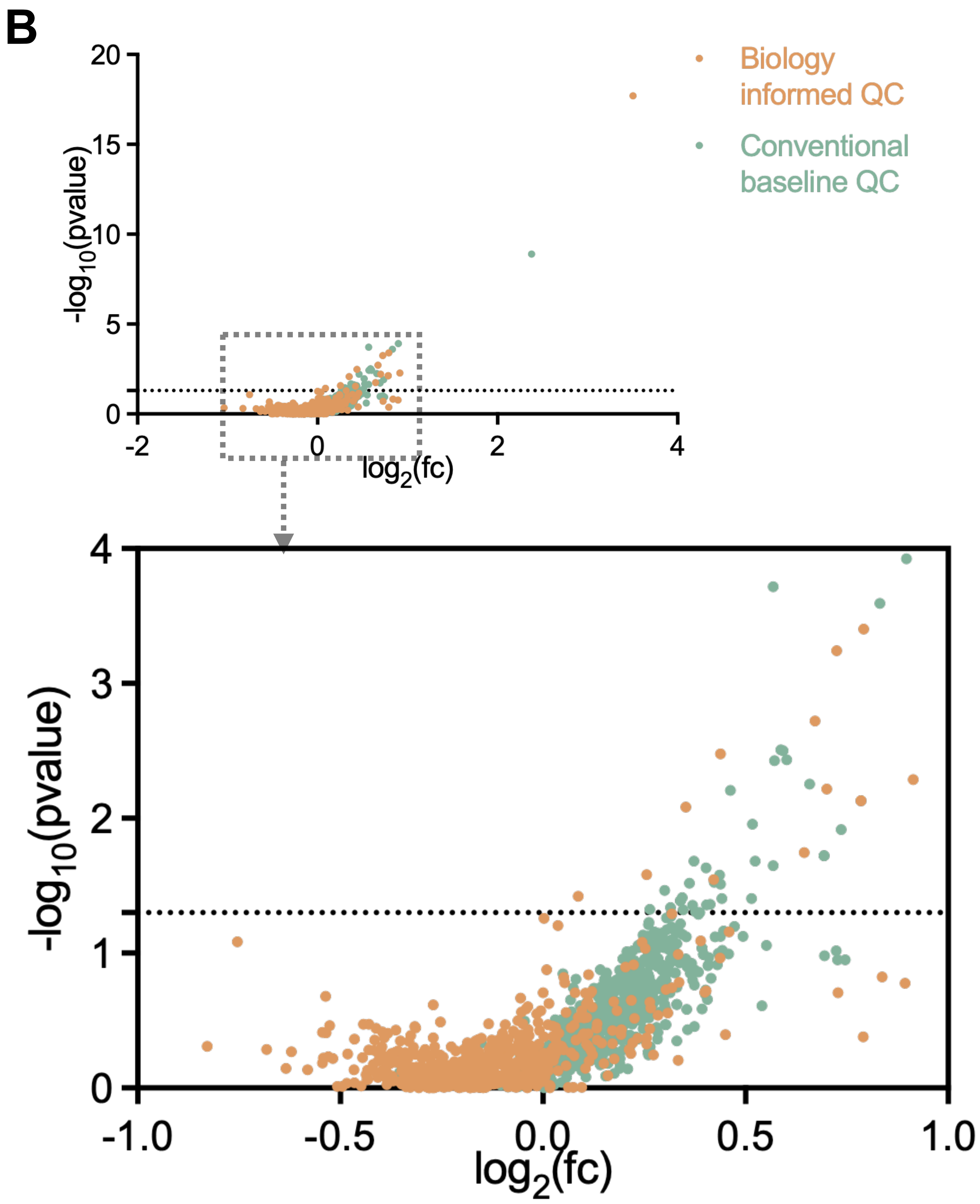


**C**

| Target | Biology informed QC | | Conventional baseline QC | |
|---|---|---|---|---|
| | $log_2$(fc) | $-log_{10}$(p-value) | $log_2$(fc) | $-log_{10}$(p-value) |
| $A\beta_{1-42}$ | 3.50 | 17.70 | 2.38 | 8.90 |
| APOE | 0.67 | 2.72 | 0.60 | 2.43 |
| C4a | 0.78 | 2.13 | 0.69 | 1.72 |
| CD11c | 0.70 | 2.22 | 0.57 | 1.65 |
| CD64 | 0.73 | 3.24 | 0.57 | 2.43 |
| EGFR (pY1068) | 0.35 | 2.08 | 0.46 | 2.21 |
| HLA-C | 0.42 | 1.55 | 0.40 | 1.36 |
| HLA-DR | 0.91 | 2.29 | 0.74 | 1.92 |
| IBA1 | 0.44 | 2.48 | 0.38 | 1.31 |
| Podoplanin/gp36 | 0.79 | 3.40 | 0.83 | 3.59 |
| SFRP1 | 0.65 | 1.75 | 0.66 | 2.25 |
| TDP-43 | 0.78 | 2.13 | 0.69 | 1.72 |

Supplementary Figure 6
