## Supplementary figures and images for "Spatial Multi-Omics Workflow and Analytical Guidelines for Alzheimer’s Neuropathology"

### Supplemental figure 4

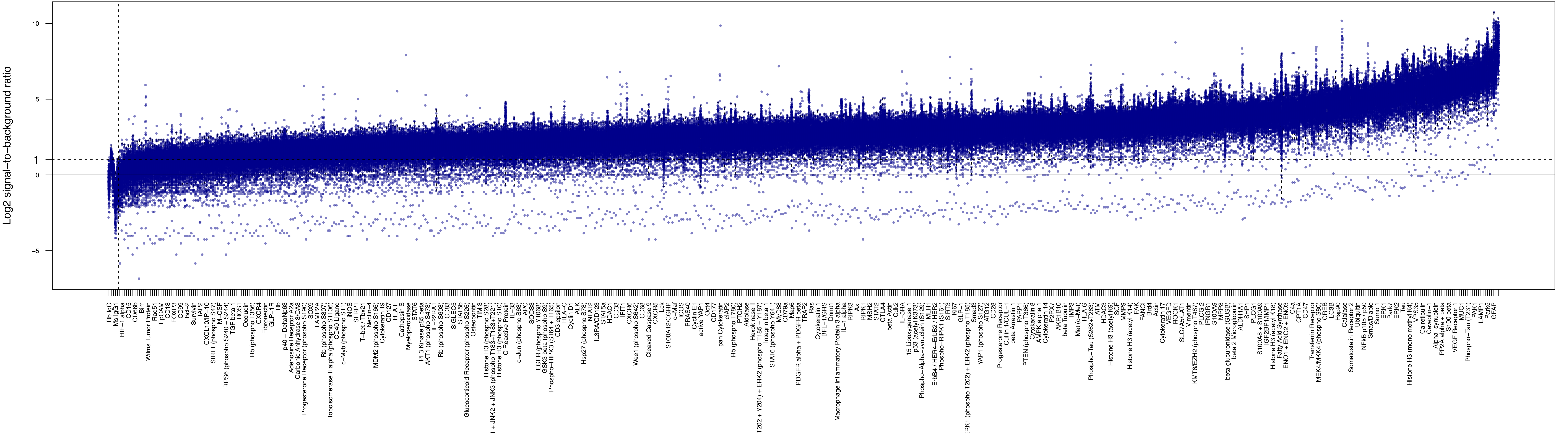
